# Bold zebrafish (*danio rerio*) learn faster in an associative learning task

**DOI:** 10.1101/2024.01.30.578005

**Authors:** Jamie Corcoran, Levi Storks, Ryan Y. Wong

## Abstract

Animals differ in their ability to learn. One potential factor contributing to learning differences is personality types. We investigated the relationship between learning and the bold-shy continuum by comparing performance of bold and shy zebrafish in conditioned place preference (CPP) and 2 choice tasks. Bold fish learned significantly faster than the shy fish but there were no differences in their final performance. When tested in the 2 choice task, we found no clear evidence of learning, however bold fish made more initial choices than shy fish. Overall,our study suggests that bold fish tend to be faster learners when compared to shy fish. The lack of differences in the final change in behavior suggests that the learning difference is due to neophobic tendencies and resulting initial interactions with the learning stimulus.

## Introduction

As animals interact with their environment, how quickly they learn and recall these interactions can vary between individuals (Boogert et al., 2018, Cauchoix et al., 2018). It has been hypothesized that variation in learning between individuals can be explained in part by differing personality types (Dingemanse & Wolf, 2010, Sih & Del Guidice, 2012, Sih et al., 2004). Across many animal taxa, studies demonstrate that one common dimension of personality is the bold-shy continuum (Réale et al., 2007). Bold individuals are characterized by displaying lower neophobic and stress-related behaviors and have higher exploratory activity. In contrast shy individuals tend to have opposing traits (Wilson et al., 1994, Sih et al., 2004, Baker et al, 2018).

However, studies across taxa find a conflicting relationship between personality and learning. Many studies showed that bold individuals learn faster than shy, in animals such as mammals, birds, to teleost fish (Mazza et al., 2018, Guenther et al., 2014, Dugatkin & Alfieri, 2002, DePasquale et al., 2014, Bensky et al., 2017, Daniel & Bhat, 2020, Kareklas, Elwood & Holland, 2017). Fewer studies either found the opposite (e.g. shy learn faster than bold) or no relationship between personality and learning speed (Lermite, Peneaux & Griffin, 2016, Ferron et al., 2015, Sommer-Trembo & Plath, 2018, Baker & Wong, 2019). Inconsistencies across studies suggest that other factors likely influence learning performance beyond personality type. Aspects of the learning assay like the type of task (e.g. operant or classical conditioning) or the context that the animal is tested in could affect the relationship between personality and learning (Poirer et al., 2020, Dingemanse & Wolf, 2010).

Individuals of varying personality types likely vary in their interactions with different learning tasks or stimuli, which may influence learning performance (Sih & Guidice, 2012). For example, different training paradigms require that the animal engage with the stimulus in different ways. Some studies found that learning is not correlated across training paradigms (Guillette et al., 2015, Ducatez et al., 2014, Kassai et al., 2022, Poirer et al., 2020) and one found that changing the difficulty of the learning task changed the relationship between personality and learning speed (Chang et al., 2018). Similarly, a meta-analysis in non-human animals found a low correlation between learning ability across cognitive tasks (Poirer et al., 2020). This potential variation across tasks suggests a need for measurements in multiple learning tasks (Griffin et al., 2015). Neophobia, associated with a shy personality, has been seen to affect operant learning of a food reward due to higher latencies to approach (Stöwe et al., 2006). Thus, comparing a passive (classical) task that does not require the animal to approach a novel object to an active (operant) task that does require approach may produce different results.

In this study, we investigated the effect of personality type on learning performance across two associative learning paradigms in zebrafish (*Danio rerio*). Using a within-subjects and counter-balanced design we individually trained bold and shy zebrafish to associate a visual stimulus with a food reward in both conditioned place preference and 2 choice tasks. We tested the prediction that bold individuals will be faster learners compared to shy fish because of their decreased neophobia. We also evaluated the prediction that there will be an interaction effect of personality and training paradigm on learning speed. Given that the operant task requires fish to actively make a choice, we expected that bold fish would learn faster in this task than shy fish due to their decreased neophobia.

## Methods

### Animals

We used zebrafish from selectively bred lines that exhibit shy (high stationary behavior, HSB) or bold (low stationary behavior, LSB) personality traits (n = 48 per line). Across six different stress and anxiety-like behavioral assays, the HSB line exhibits a greater amount of behaviors consistent with a shy personality type (e.g., freezing, less exploratory, higher cortisol levels) than the LSB line (Wong et al., 2012, Baker & Wong, 2019). Additionally, the exploratory behavior of the lines in an open field test is repeatable and reliable (Baker & Wong, 2019). The HSB line also shows faster release of cortisol under stress compared to the LSB line (Wong et al., 2019). For simplicity, we will refer to the HSB and LSB lines as shy and bold personality types, respectively. The fish used in this study were selectively bred for 13 generations from wild caught zebrafish. Before testing, we housed the fish together in 40L tanks and fish were fed twice a day with Tetramin Tropical Flakes (Tetra, USA). One week prior to testing we physically isolated fish into 3-liter tanks on a recirculating water system (Pentair Aquatic Eco-Systems or Aquaneering) using UV and solid filtration on a 14:10 L/D cycle at a temperature of 27 °C. Fish had visual and olfactory access to each other. Starting three days before testing we withheld food from the fish to reduce the possibility of satiation while training.

### Behavioral assays overview

We conducted four behavioral assays on each fish: an open field test (OFT), a test for food motivation, a 2 choice discrimination task (operant conditioning), and a conditioned place preference (CPP, classical conditioning) task. The OFT and food motivation test were performed prior to training. Using a within-subjects design, we tested each fish in both associative learning paradigms and counterbalanced the starting paradigm (Figure 1). We used frozen adult brine shrimp (*Artemia* spp., San Francisco Bay Brand, USA) administered in liquid form as the food reward. Half of the fish received distilled water instead of brine shrimp to serve as controls. We started with four groups of 24 fish (bold control, shy control, bold treatment, and shy treatment).All behavioral assays were performed between 3-8 hours after light onset. After 4 days of isolation, we tested each fish in the open field test to validate behavioral phenotype. We then habituated each fish for two consecutive days in the conditioning tank. We assessed biases in food motivation for the brine shrimp before starting baseline trials of the associative learning assays. Each fish had a 14 day inter-assay testing interval to minimize the influence of the tasks on each other.

**Figure 1.**
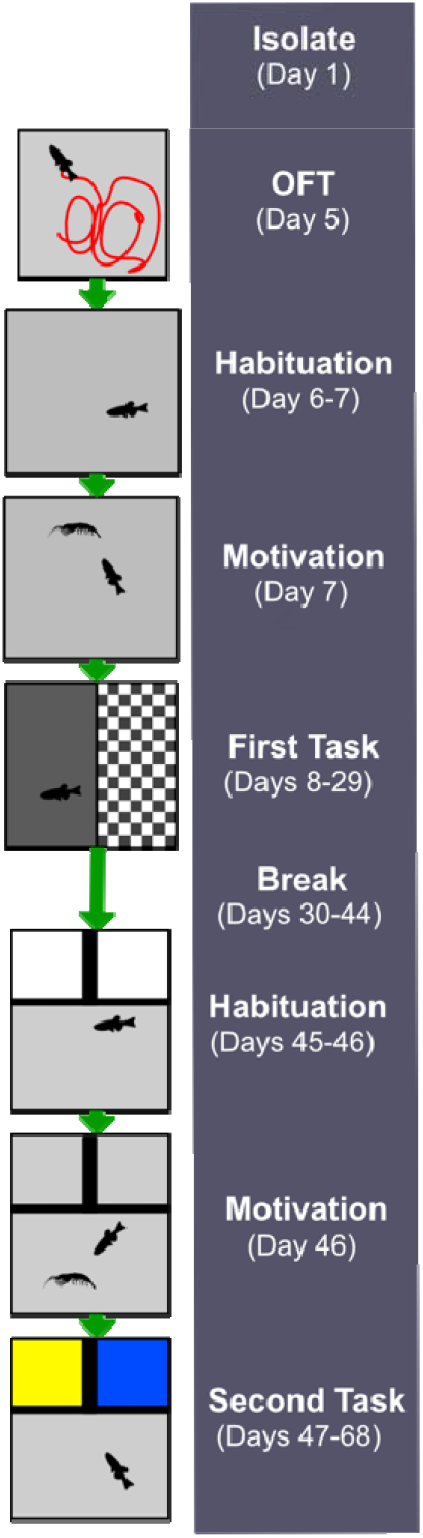
Overview of Experiment Timeline. Fish of all groups started with isolation on the first day then went through an open field test (OFT), habituation, motivation test, and training on the first task (conditioned place preference task in this illustration). After a break fish went through the same for the second task (2 choice discrimination task in this illustration).Our study design was counterbalanced and half of the fish began with the CPP task while the other half began with the 2 choice discrimination task.

### Open field test

We individually tested fish in an OFT in a tank that was 31.75cm x 31.75cm x 10cm containing 4L of water. Immediately after placing fish in the tank we video-recorded the individual’s behaviors for 5 minutes. We used Ethovision XT 17 (Noldus, Netherlands) to quantify the amount of time that each individual spent frozen during the trial.

### Motivation test

This test was performed in the AD and LT models of the Zantiks semi-automated behavioral units (Zantiks, Cambridge, UK). After 30 seconds for acclimation, the food reward was administered 3 times at 30 second intervals. We quantified the time spent in a 9x12 cm rectangle centered around the food administration tube. The time that was being measured started immediately after the first brine shrimp administration until the end of the test to measure the motivation of the fish for the food reward. We performed the test in both Zantiks models but due to the size and height of the tank in the larger LT unit, the food drifted outside the fish tracking zone. Thus, we only used the data from the AD unit to assess motivation.

### Conditioned place preference

We used a modified conditioned place preference protocol (Lau et al., 2006) in the Zantiks LT unit. The testing tank (36 cm x 27 cm x 30 cm) was filled with 5.8 L of water. We tested each fish in the CPP task for three weeks that consisted of 2 days of habituation, 1 day of baseline testing, 11 days of conditioning, and 3 days of probe trials (Figure S1a). To habituate each fish to the assay we placed the fish in the tank for 10 minutes with no training stimulus lights. After habituation we determined the baseline preference for the light stimuli (gray or checkered pattern) for each fish. Fish swam freely for 10 minutes in the tank where one half was illuminated from the bottom with a gray screen and the other half a checkered screen. We determined the conditioned and non-conditioned stimuli as the stimulus where the fish spent the least and most amount of time, respectively. During conditioning days, we sequentially presented each stimulus for 5 minutes to each fish. The non-conditioned stimulus was presented for the first five minutes followed by the conditioned stimulus. One hundred microliters of brine shrimp or distilled water was administered every minute during presentation of the conditioning and non-conditioning stimulus, respectively. Food reward consisted of 11.4 grams of frozen brine shrimp in 30 mL of distilled water. We fed control fish an equivalent amount of brine shrimp after each conditioning trial. Probe trials were administered the day after a conditioning trial. Probe trials were conducted after 3 days, 7 days, and 11 days after the first day of conditioning with a total of 3 probe trials. Probe trial methods were the same as those used in the baseline preference step where we quantified the time spent in each stimulus for each fish. The order of stimulus presentation was consistent within a fish but random across fish for probe and baseline trials.

### 2 choice discrimination task

We used a modified 2 choice discrimination task from an established protocol (Bilotta et al., 2006). We used the AD model Zantiks unit (Zantiks, Cambridge, UK) with a 14cm x 20cm x 15cm tank filled with 2.5 L of water (Figure S1b). We habituated each fish for 20 minutes a day for two consecutive days with white lights on in the wells as shown in Figure S1b. We tested each fish every other day for a total of 10 testing days. Fish were fasted on non-testing days. In this task the fish were presented with two 6.5 cm x 5.1 cm light stimuli (blue and yellow) from below at one end of the tank. Prior studies show that with appetitive learning in zebrafish there is a bias towards red compared to other colors such as blue and yellow (Spence & Smith, 2008, Kim et al., 2017). For each fish, a color was randomly chosen at the start of testing to be the reinforced stimulus where a food reward (brine shrimp) was administered at the other end of the tank when the fish swam into the designated reinforced color. The food reward consisted of 5.7 grams of frozen brine shrimp suspended in 30 mL of distilled water. Each trial began with an acclimation period of two minutes with white lights in the two wells. After two minutes blue and yellow lights were presented for 30 seconds. Swimming into the designated correct choice resulted in the correct colored light staying on for an additional 30 seconds and we simultaneously administered 25 _μ_l of the food reward. An incorrect choice resulted in all lights turning off for 30 seconds. This sequence ran for a total of 20 trials each day for each fish (i.e., one session consists of 20 trials). The position of the yellow and blue lights (e.g., left or right) was randomly set for each trial. There was an intertrial interval of 10 seconds. Control fish underwent the same protocol with distilled water administered instead of brine shrimp and were fed brine shrimp after each testing day. We compared the number of correct choices and the total number of choices across sessions to assess learning.

### Statistical analysis

We performed all statistical tests using R statistical software(R 4.2.2 GUI 1.79 Big Sur ARM build) and Rstudio version 2022.12.0+353 (R Core Team, 2021). Due to fish mortality during the experiment, the sample sizes for statistical analyses between the conditioned place preference (bold control (n = 20), shy control (n = 20), bold treatment (n = 19), and shy treatment (n = 19)) and 2 choice (bold control (n = 20), shy control (n = 17), bold treatment (n = 19), and shy treatment (n = 20)) tasks differed. We conducted post-hoc tests using the emmeans (Lenth et al., 2022) package and normality and assumptions were checked using base R. The lme4 package (Bates et al., 2022) was used to test negative binomial linear mixed effect models. We obtained simple statistics for all measures using the psych package (Revelle, 2022) (Table 1). Sex was included in all models but was not significant and therefore removed. Model assumptions, including normality were inspected in R.

### Open field test and motivation

We tested for differences between the bold and shy groups in the OFT and motivation test using a Welch two-sample t-test. This test was used due to unequal variances between bold and shy groups. We compared the duration of time frozen in the OFT between the bold and shy personality types. To investigate difference in food motivation, we compared the duration of time spent around the food administration tube between the bold and shy personality types using the same test.

### Conditioned place preference

We modeled the duration of time spent in the conditioned stimulus for the last half (5 minutes) of the baseline and probe trials to test for a change in preference for the conditioned stimulus across the task within the different groups. We did not include the first half (5 minutes) in the analysis to minimize the influence of handling on fish behavior. We performed a repeated measures ANOVA to investigate the effects of treatment, personality type, and conditioning day on the time spent in the conditioned stimulus with a linear mixed effects model with individual as the random effect. We included all interactions in the model and used type II sums of squares. We used Tukey post-hoc tests to evaluate differences in the response variables across trials for each group and within trials between groups.

### 2 choice discrimination task

We modeled the number of correct choices over the conditioning days to examine changes in correct choices over time within groups. We performed a negative binomial mixed effect regression on the number of correct choices with treatment, personality type and session as the fixed effects and ID as the random effect. Simple slopes were obtained to test for increases in correct choices within each group using the interactions package in R and plotted using the same package. Additionally, we performed a negative binomial mixed effect regression on the total number of choices with treatment, personality type and session as the fixed effects and ID as the random effect. We also obtained simple slopes for this model.

## Results

### Shy fish freeze more but had equal motivation to eat

There was a significant effect of personality type on freezing time in the open field test (Figure 2a). Shy fish spent significantly more time frozen than bold fish (t =-3.55, df = 90, p = 6.4*10^−4^). There were no significant differences between personality types (t = -0.19, df = 82, p =.85) in the amount of time spent around the food in the motivation task (Figure 2b).

**Figure 2.**
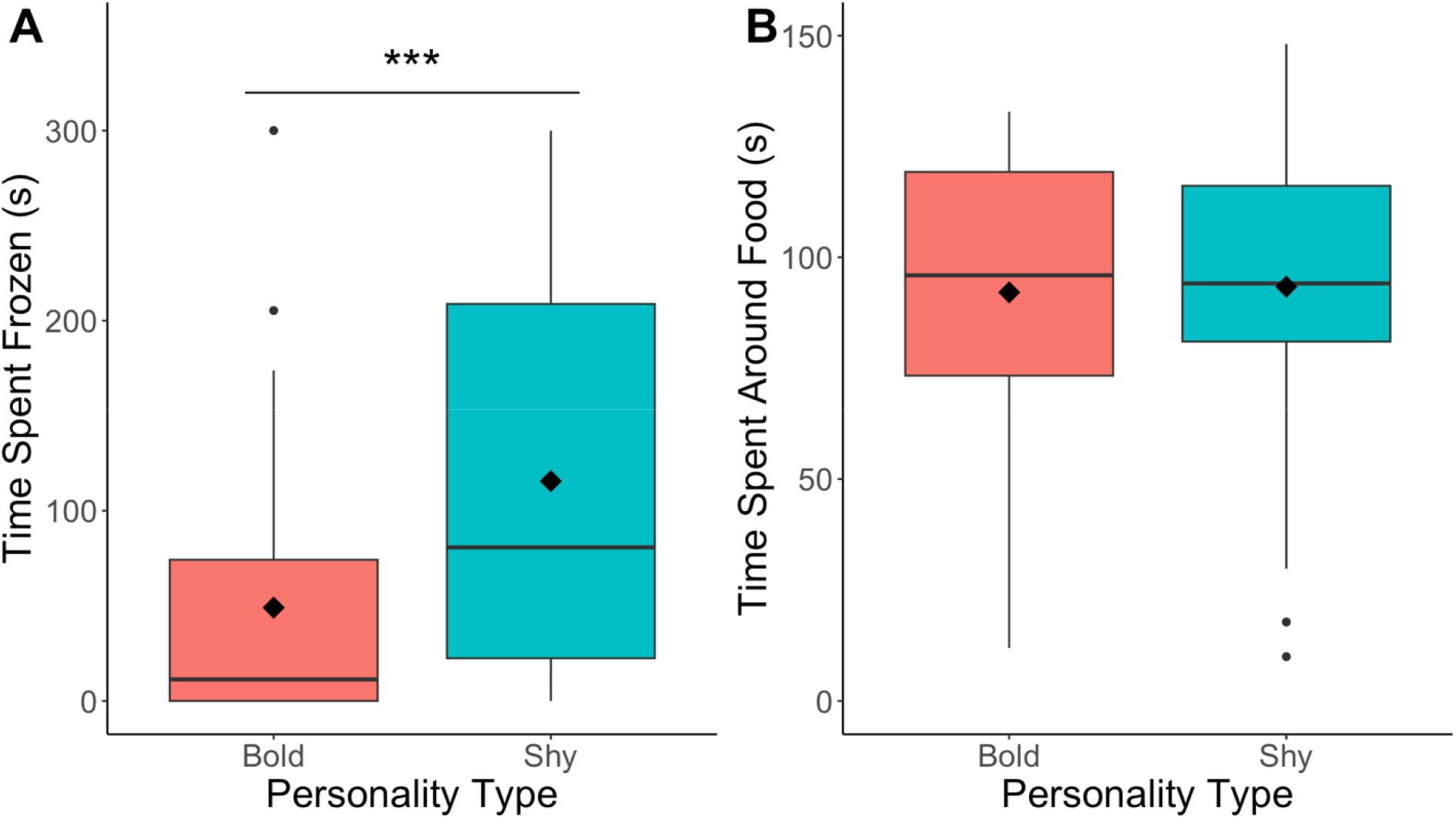
A. Boxplot of time spent frozen in the open field test and B. boxplot of time spent around the food in the motivation task. Bold fish are in red and shy fish are in teal. The diamond indicates the mean and the line is at the median.*p<.05, ** p <.01 ***p<.001

### Bold fish change their behavior before shy fish

Treatment fish increased time spent in the conditioned stimulus in the CPP task, with bold fish increasing time spent in the conditioned stimulus earlier in the task than shy fish (Figure 3). In the full model (Table S1) the interaction effect between treatment and probe trial was approaching significance (F(3, 292) = 4.09, *p* = .09). A Tukey post hoc test (Table S2) revealed that there were no significant differences in the duration of time in the conditioned stimulus between trials for the control groups for either personality type (*p* > .05). In the bold treatment group there was a significant difference between baseline and probe 1 (i.e., after 3 days of conditioning; t = -2.64, df = 296, *p* = .03), probe 2 (i.e., after 7 days of conditioning; t = -3.55, df = 296, *p* = 2.8*10^−3^), and probe 3 (i.e., after 11 days of conditioning; t = -3.35, df = 296, *p* = 4.7*10^−3^). In the shy treatment group there was no significant difference in time spent in the conditioned stimulus between baseline and probe 1 (t = -1.27, df = 296, *p* = .45) but there was a trend for a difference between baseline and probe 2 (t = -2.42, df = 296, *p* = .07) and at probe 3 shy treatment group spent significantly more time in the CS compared to baseline (t = -2.67, df = 296, *p* = .04). No significant differences in duration of time in the CS between probe 1, 2, or 3 were detected in any of the groups (*p* > .05). There were no differences in time spent in the conditioned zone at any of the time points between personality types (*p* > .05). Additionally, there was no significant correlation between learning speed (change in CS time from baseline after 3 days of conditioning) and final time spent in the conditioned stimulus in the CPP task for the bold fish (_ρ_ = .19, *p* = .44) or for the shy fish (_ρ_ = .22, *p* = .35).

**Fig. 3.**
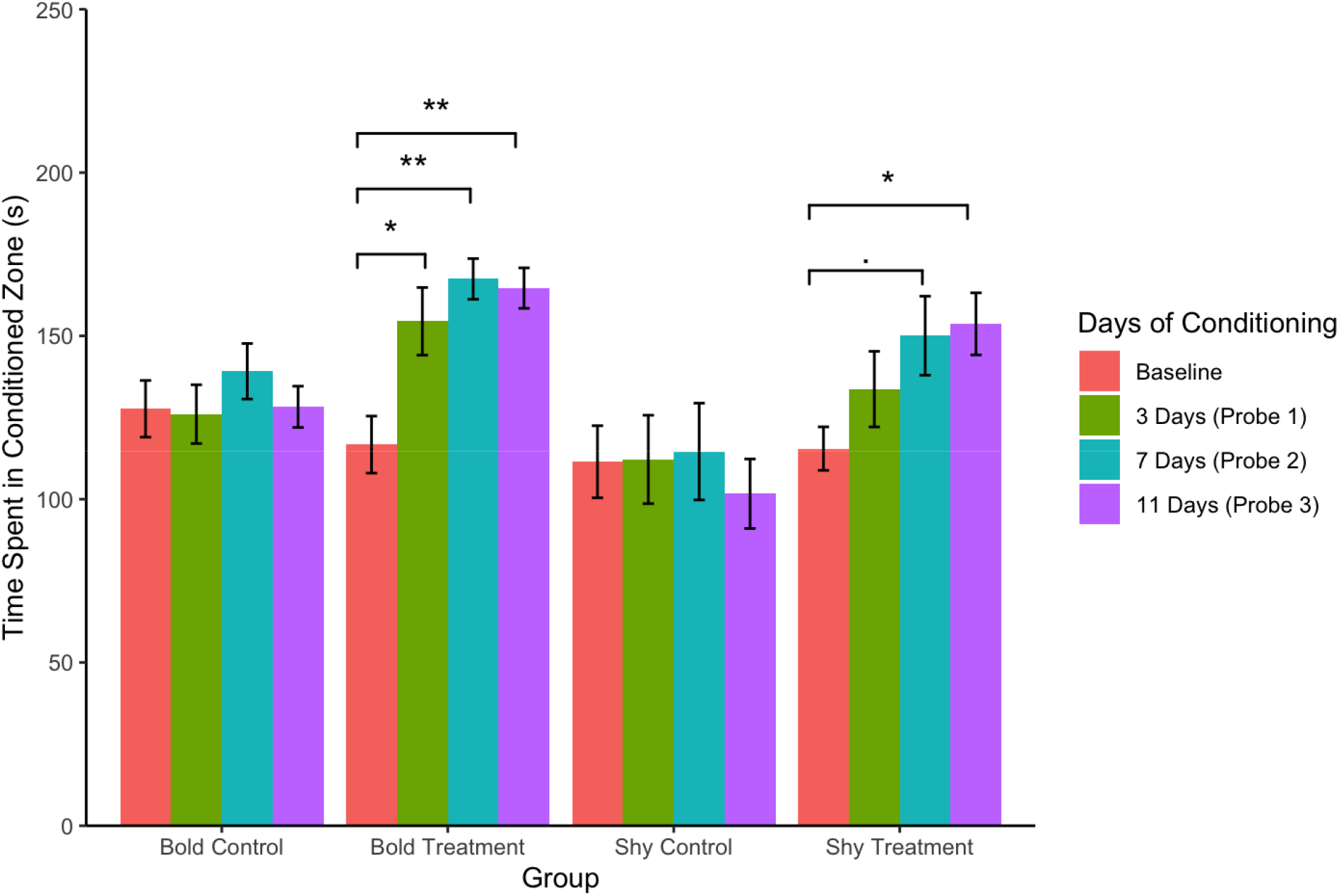
Time spent in the conditioned zone by group and day of conditioning in the CPP. Pink bars are at baseline, green bars are after 3 days of conditioning (Probe 1), blue bars are after 7 days (Probe 2) and purple bars are after 11 days of conditioning (Probe 3). Error bars indicate standard error. .*p* < .1, \**p* < .05, \*\**p* < .01.

### No evidence of learning in 2 choice discrimination task with correct choices

In the 2 choice discrimination task there was no significant difference in number of correct choices between control and treatment fish (Table S3). There was only a significant main effect of personality type such that bold fish made more correct choice compared to shy fish (b = -.49, t = -2.84, *p* = .01) and a significant interaction between personality type and session (b = 0.03, t = 2.601, *p* = .01). Testing for the simple slopes (Table S4, Figure S2), both shy control (m = 0.03, t = 3.15, *p* = 2.2*10^−5^) and shy treatment (m = 0.04, t = 4.24 *p* = 4.4*10^−6^) groups had a significant positive slope while bold control (m = 0, t = -0.26, *p* = .79) and bold treatment (m = 0.01, t = 0.98, *p* = .33) have no significant relationship.

### Difference across treatment and control only in total number of choices in 2 choice discrimination task

For the total number of choices, there was a significant difference between control and treatment fish (Figure 4, Table S5). There was a main effect of personality type on total number of choices (b = -.34, t = -2.07, *p* = .04) where bold fish had higher total number of choices than shy. The interaction between session and treatment is approaching significance (b = 0.15, t = 1.69, *p* = .09). Testing for the simple slopes, shy control (m = 0.01, t = 1.22, *p* = .22), and bold control (m = 0, t = -0.44, *p* = .66) did not have a significant relationship (Figure 4a). Only shy treatment (m = 0.03, t = 4.59, *p* = .4.4*10-6) had a significant positive slope (Figure 4b, Table S6). In contrast, bold treatment had a slope approaching significance (m = 0.01, t = 1.90, *p* = .06) (Figure 4b).

**Figure 4.**
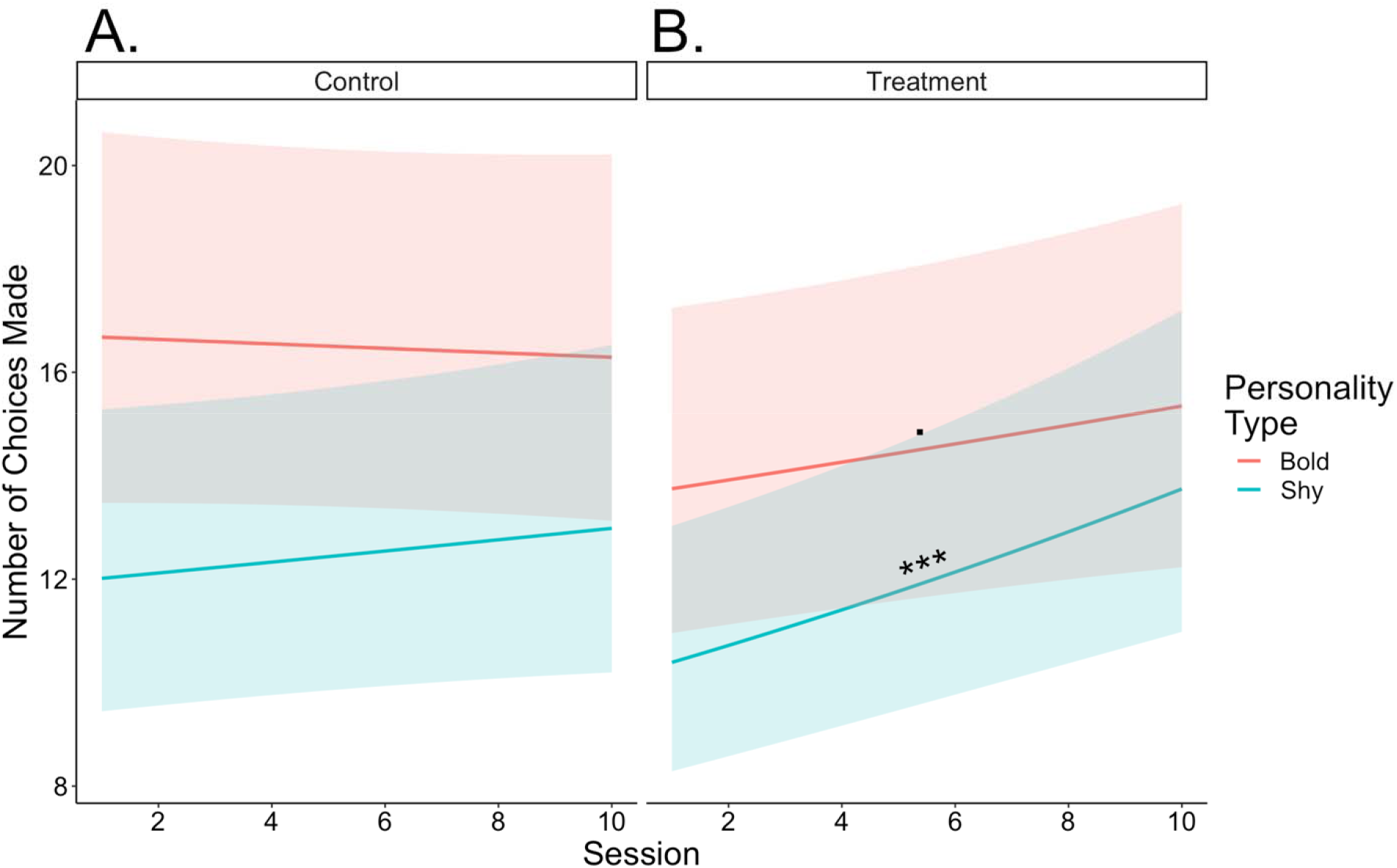
Regression lines of the number of choices made by personality type and treatment. 4A. shows the regression lines for control fish and 4B. shows regression lines for treatment fish. The bold group is in red and the shy group is in blue. Shaded regions indicate a 95% confidence interval. The simple slopes significance is indicated, *p* < .1, \**p* < .05, \*\**p* < .01, \*\*\**p* < .001.

## Discussion

Variation in learning performance can be due to complex interactions between intrinsic (e.g., personality type) and extrinsic (e.g. learning task) factors (Sih & Guidice, 2012). We investigated the effects of personality type and learning task by testing zebrafish of differing personalities across two different associative learning assays. Overall, we found that learning performance in one of the tasks was influenced by an animal’s personality type.

Bold fish increased time spent in the conditioned stimulus earlier than shy fish in the conditioned place preference task, which suggests that bold fish learned faster in this task. The bold fish showed a significant increase in time spent in the conditioned stimulus after just 3 conditioning days whereas shy fish took an additional 8 days of conditioning to show a significant change from baseline (Figure 3). These results are consistent with other studies demonstrating that individuals with bold personality types learn faster than shy individuals (Mazza et al., 2018, Guenther et al., 2014, Dugatkin & Alfieri, 2002, DePasquale et al., 2014, Bensky et al., 2017, Daniel & Bhat, 2020, Kareklas, Elwood & Holland, 2017). Differences in learning speed between personality types in this task may be due to differences in behavior such as stress reactivity, exploration, and neophobia (Sih & Guidice, 2012, Sommer-Trembo & Plath, 2018). Our observed differences in learning speeds between personality types cannot be explained by differences in motivation for the food reward (Figure 2). Interestingly, there were no differences in the amount of time spent in the conditioned stimulus between the personality types after 11 days of conditioning, suggesting that individuals approach an asymptotic level of performance. This suggests that both personality types are capable of changing their behavior (e.g. learn) to similar extents and therefore differences in cognitive ability between personality types is an unlikely explanation for differences in learning speed.

When testing the same fish in the 2 choice discrimination assay, there was no significant difference in the number of correct choices between treatment and control groups, which suggests the fish did not learn the stimulus-reward association in this task (Figure 4). However, there were differences across personality types in which both shy treatment and control increased the number of correct choices while the bold groups did not. The positive slope for the shy groups is likely due to an overall increase in total choices with repeated exposure. When looking at the total number of choices made over sessions, the control groups did not change over time while the treatment groups increased the total number of choices made over sessions. This suggests that the treatment fish did not learn the color association but may instead have learned to go into the wells. Animals can attend to several cues in discrimination learning and sometimes attend to unintentional or general cues (Mackintosh, N., J., 1965). We also cannot rule out that rewarding the fish in a different location than the stimuli may have decreased the strength of pairing between action and reward (Murphy & Miller, 1958). While in the 2 choice discrimination task fish did not learn the color association, the bold fish made more choices than shy fish in the first session. This is likely due to decreased neophobia and increased exploration in the bold fish as demonstrated in the open-field test (Sih et al., 2004, Wong et al., 2012).

Differences in neophobia (e.g. latency to approach novel objects) classically distinguish bold and shy personality types (Carter et al., 2012, Sih et al., 2004, Wilson et al., 1994). In the current study one potential explanation for bold fish learning quicker in the conditioned place preference and making more initial choices in the 2 choice discrimination task relative to shy fish are differences in neophobia between the personality types. The shy fish could have found the colored lights in the 2 choice discrimination task initially aversive and increased their choices as they habituated to the novel stimuli. Shy individuals tend to have increased neophobia and habituate slower, which would result in the shy fish taking longer to make active choices (Carter et al., 2012). The two days of habituation in the 2 choice discrimination task only allowed the fish to experience the tank and lighted wells but at start of conditioning they were naïve to the color of the lights and the changing stimuli. A similar effect was seen in *Gallus gallus* where individuals that were less exploratory (i.e., shy) habituated slower to a loud sound than those that were more exploratory (Dissegna et al., 2022). Neophobia may also explain shy fish learning slower in the CPP task, as shy fish could have experienced more stress than the bold fish at the start of the task even after habituation and so learned the positive association slower. Mollies (*Poecilia mexicana*) that were desensitized to the lights and sounds used in the task showed no differences in learning related to personality type (Sommer-Trembo & Plath, 2018). Increasing familiarity with the task environment and stimuli could explain why shy fish were slower to increase their preference for the conditioned stimulus but ultimately reached a level of performance similar to bold fish after 11 days of conditioning. Bold individuals tend to make associations faster likely because they are less neophobic and in a simple conditioned place preference task, this leads to them learning faster but does not change the plateau of performance (Dugatkin & Alfieri, 2002, DePasquale et al., 2014, Daniel & Bhat, 2020).

The relationship between more rapid learning and bold personality type is not consistent across all studies (Ferron et al., 2015, Lermite, Peneaux & Griffin, 2016). Potential explanations are that the relationship between speed of learning and personality can depend on aspects of the task such as learning stimulus valence or task complexity. Shy zebrafish trained in a contextual fear learning paradigm showed faster learning than bold zebrafish (Baker et al., 2019). As shy zebrafish have a faster glucocorticoid response to a novelty stressor than bold fish, this may facilitate quicker learning of aversive stimuli (Wong et al., 2019, Rau et al., 2005, Riggenbach et al., 2019) but inhibit learning of appetitive stimuli seen in current study. For task complexity, a study looking at learning accuracy found that aggressive spiders (e.g. bold personality type) were more accurate in a simple task but not in a more complex task (Chang et al., 2018). Future work may consider testing whether the same trend holds in a more complex classical conditioning task. In a more complex task, bold fish may make incorrect associations and not learn as quickly as shy fish.

Overall, we found support for differences between bold and shy individuals in how they interact with two different learning tasks. These differences in performance could be explained by varying neophobia between bold and shy individuals. In a 2 choice task requiring an active behavioral response, we found differences in initial number of choices made between personality types, suggesting that the personality types naively interacted with the stimulus differently. In the conditioned place preference task, the bold fish learned faster than the shy fish. Differences in performance between bold and shy individuals in both tasks could be explained by variation in neophobia related to personality type. Additionally, the bold and shy fish reached a similar level of performance. We encourage future studies to test the performance of bold and shy individuals across different tasks to compare their behavior both within and across tasks. Future work should also consider explicitly measuring how individuals interact with the task environment, perhaps measuring neophobia and motivation for the task.

## Acknowledgements

We thank Elizabeth Stone and Shar Soe for help with fish husbandry. We are grateful toBrandon Villanueva Sanchez, Vy Nguyen, Abigail Reynolds, Annabella Madsen, Maddison Thurber, Sydney Klucas for helpful discussions and technical assistance. This project was funded by the National Science Foundation (IOS-1942202 to RYW), and UNO Graduate Research and Creative Activity Grant and Rhoden Fellowship to JC.

## Notes

### Competing Interest Statement

The authors have declared no competing interest.

### Summary of Updates

Figure 3 revised to be labelled correctly. Minor revisions to wordings.

