## Supplementary Figures and Tables for "Bold zebrafish (*danio rerio*) learn faster in an associative learning task"

Journal of Animal Cognition

Jamie Corcoran^1^

Levi Storks^2,3^

Ryan Y. Wong^1,2^

^1^University of Nebraska at Omaha Psychology Department, Omaha, NE USA

^2^University of Nebraska at Omaha Biology Department, Omaha, NE USA

^3^University of Detroit Mercy Biology Department, Detroit, MI US

**Supplementary Table 1**

*ANOVA of treatment, personality type, and probe on the amount of time spent in the conditioned zone in the conditioned place preference task*

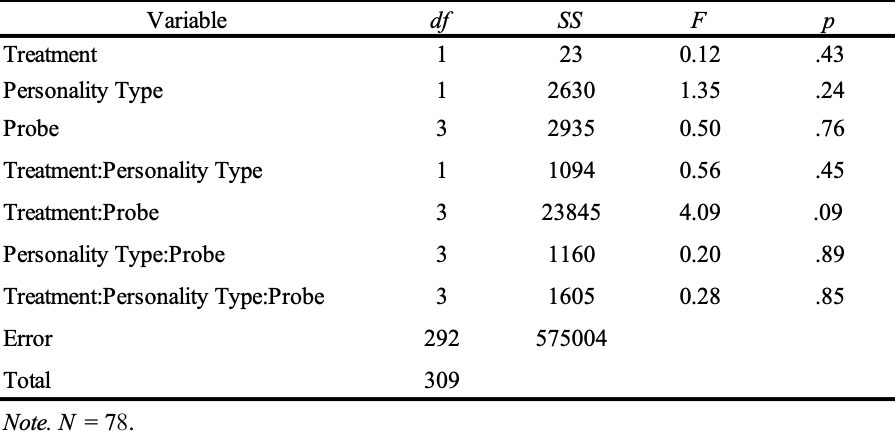

**Supplementary Table 2**

*Tukey post hoc comparing time spent in the conditioned zone at baseline, probe 1 (3 days of conditioning), probe 2 (7 days), and probe 3 (11 days) in the conditioned place preference task*

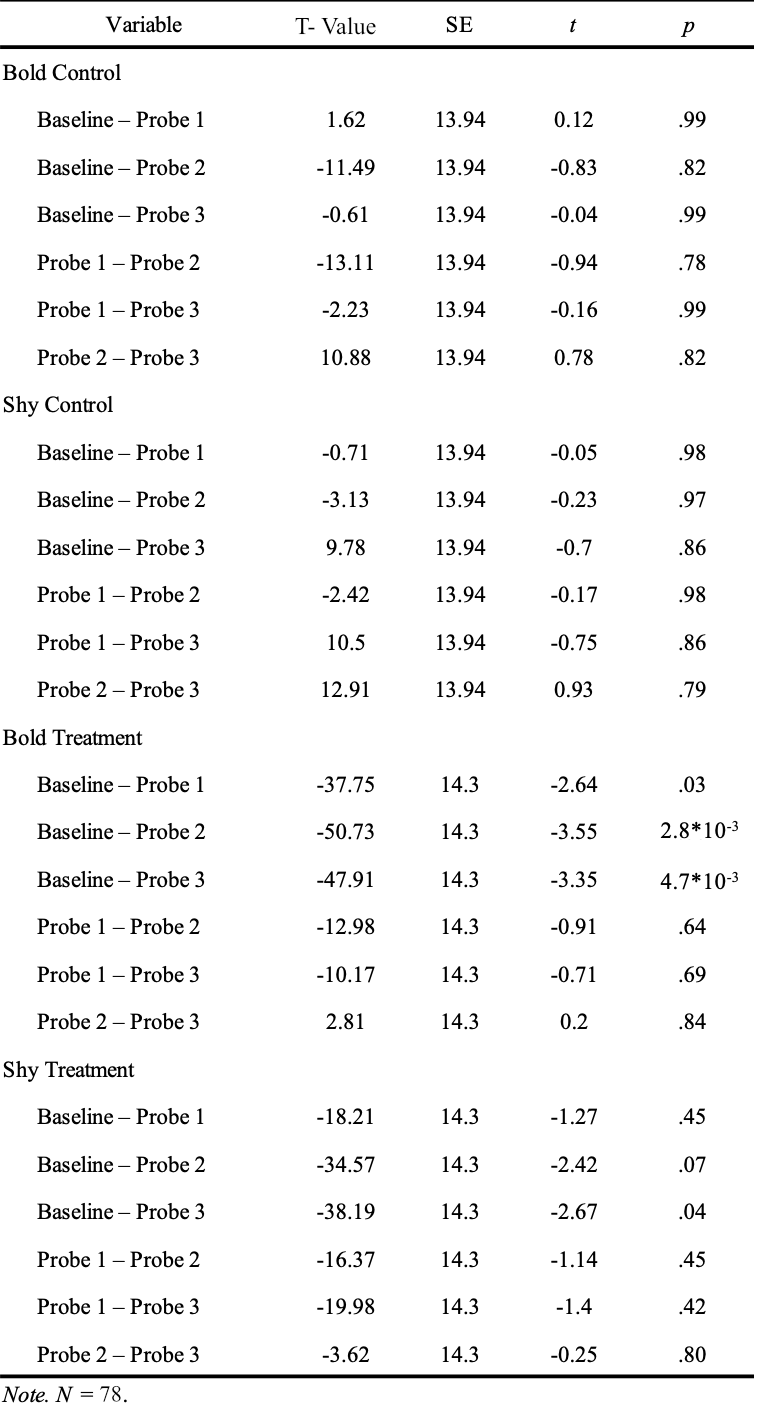

**Supplementary Table 3**

*Negative binomial mixed effect model of session, treatment, and personality type on the number of correct choices made in the 2 choice discrimination task*

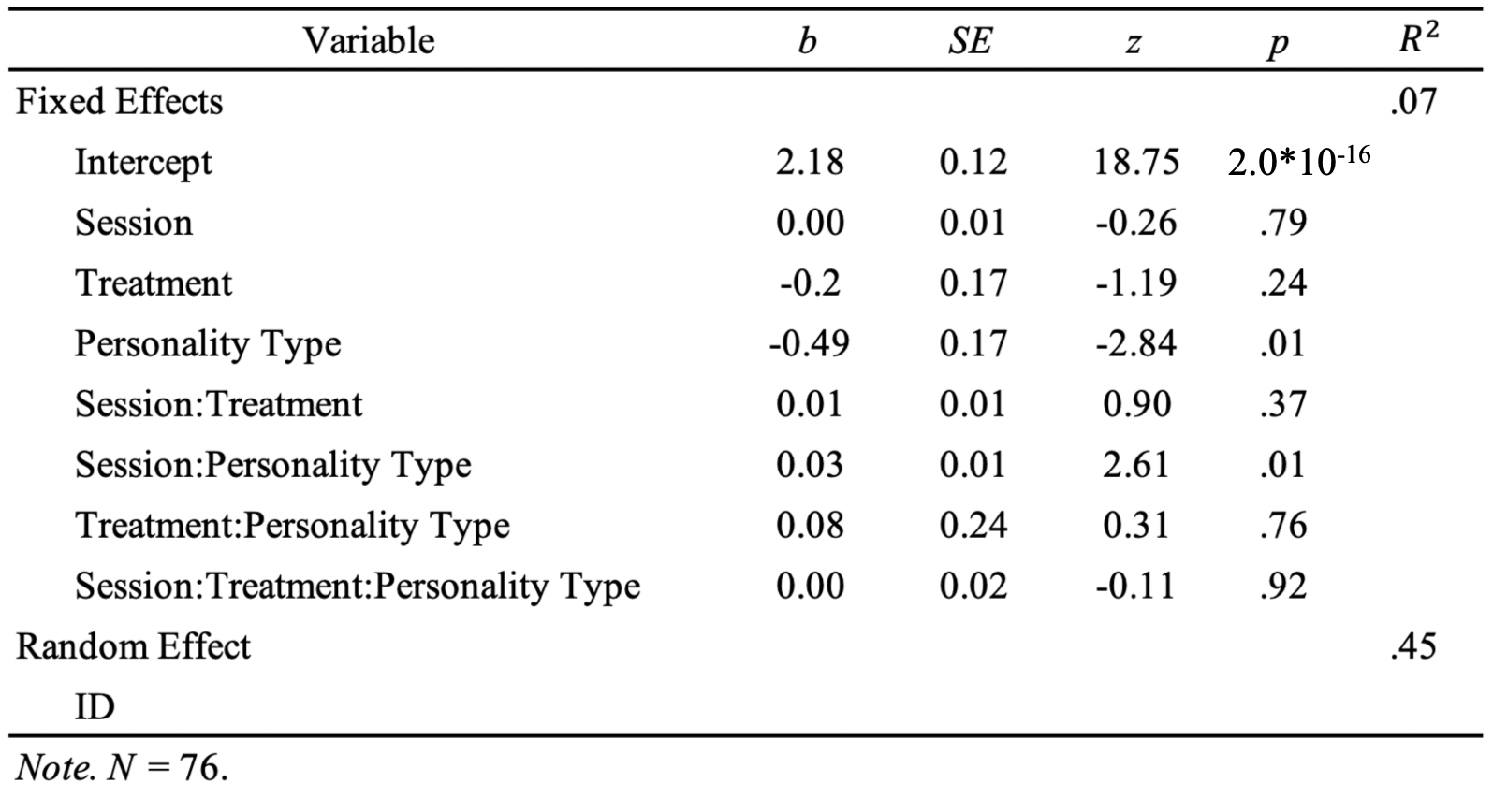

**Supplementary Table 4**

*Simple slopes of negative binomial mixed effect model of session, treatment, and personality type on the number of correct choices made in the 2 choice discrimination task*
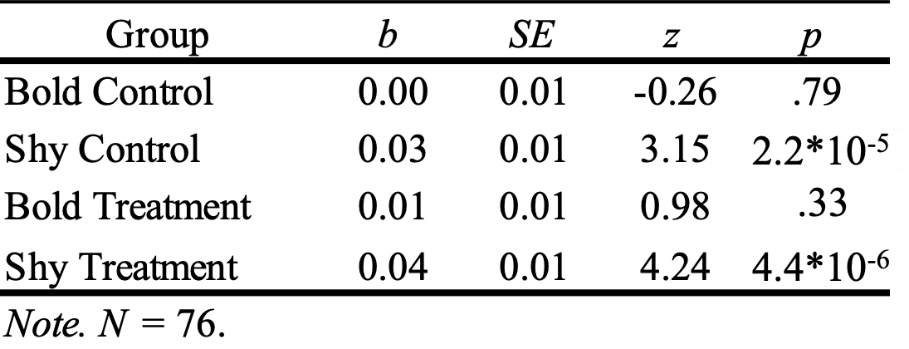

**Supplementary Table 5**

*Negative binomial mixed effect model of session, treatment, and personality type on the number of choices made in the 2 choice discrimination task*

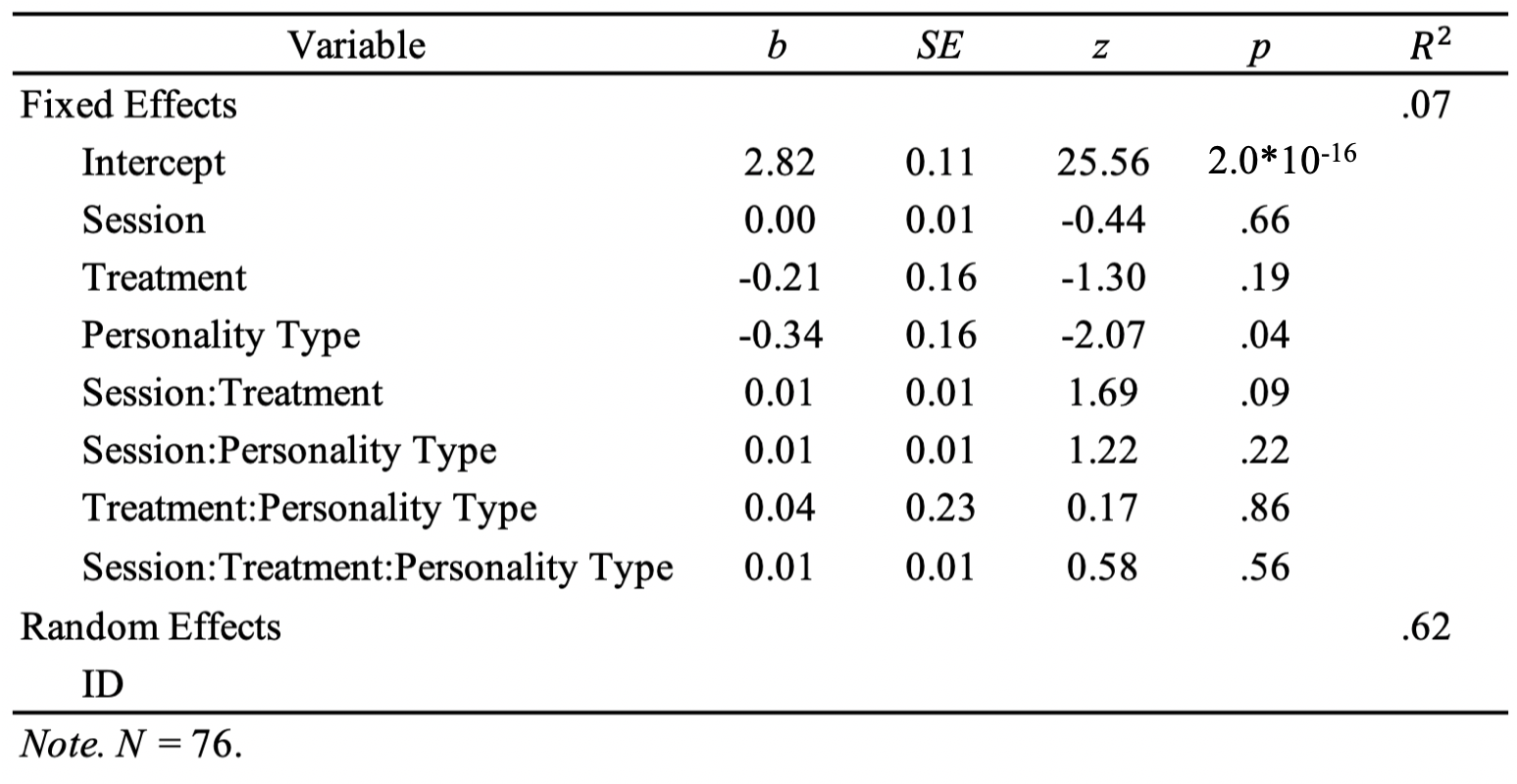

**Supplementary Table 6**

*Simple slopes of negative binomial mixed effect model of session, treatment, and personality type on the number of choices made in the 2 choice discrimination task*

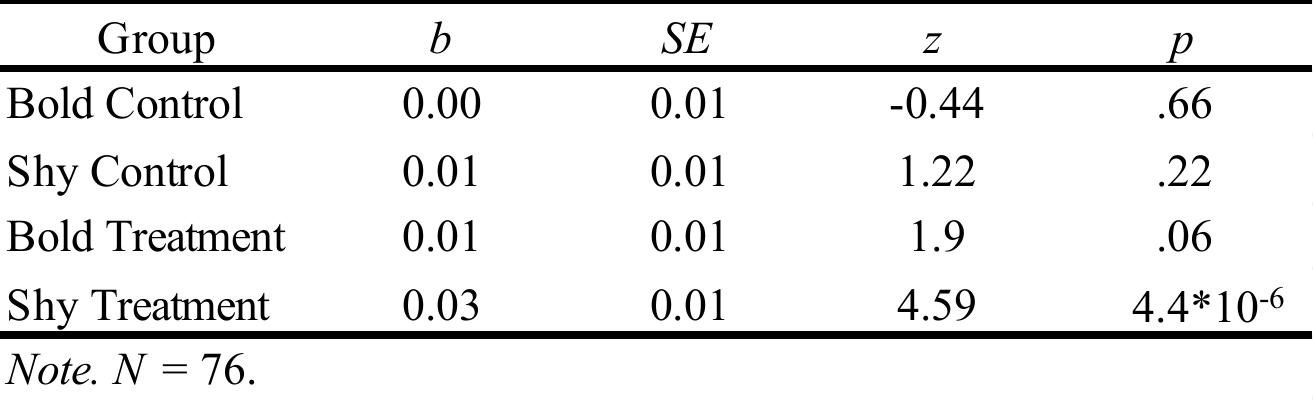

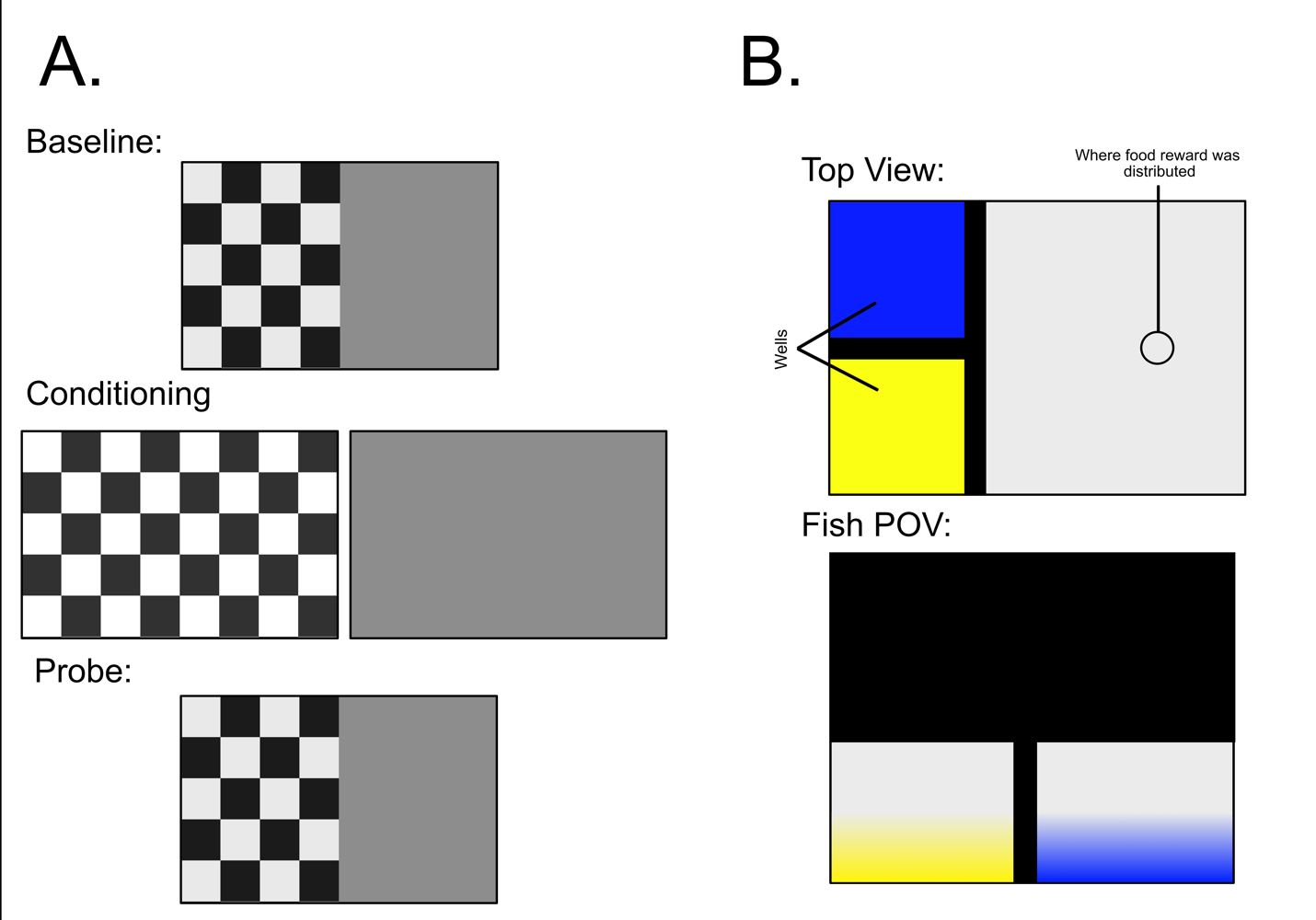

**Supplementary Figure 1.** Representation of the two tasks. (A) shows a brief overview of the stimulus displayed to the fish during the conditioned place preference task. The fish are initially presented the check and the grey stimulus. Whichever is least preferred is rewarded in conditioning trials. Then the prove presents both stimuli and the preference is measured again. B) shows the top view and the fish point of view of the 2 choice discrimination task. The wells and where the food reward was distributed are labelled.

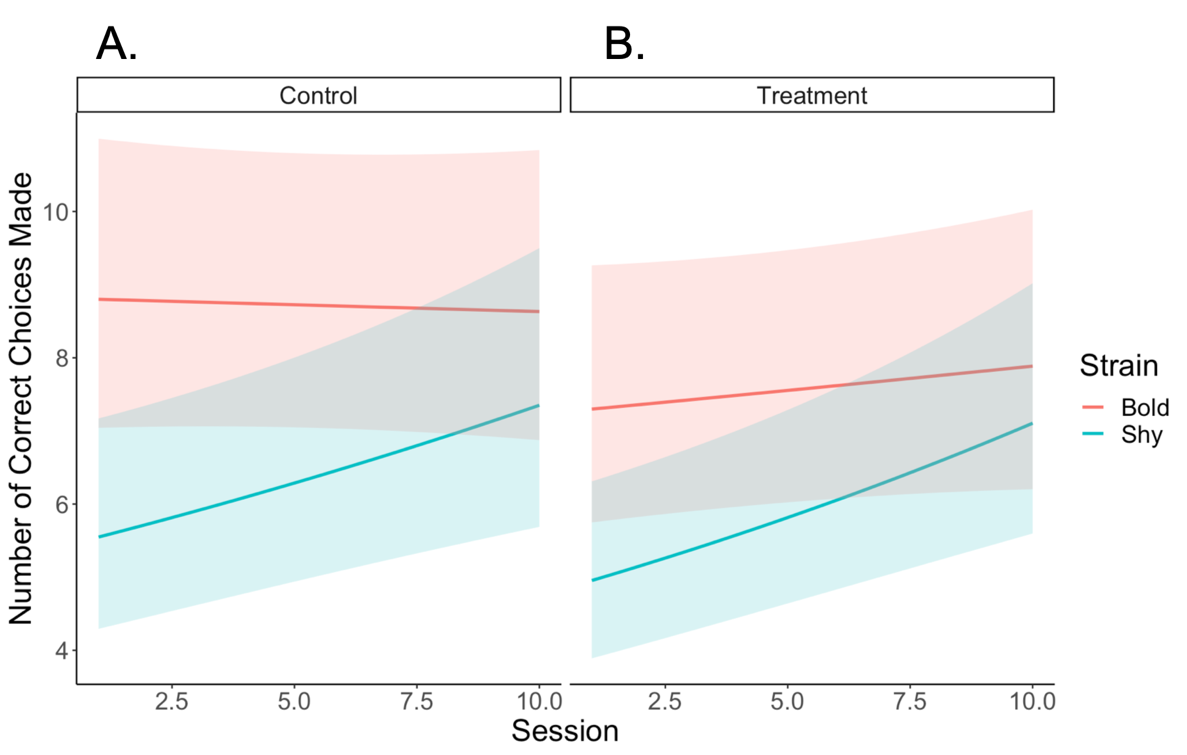

**Supplementary Figure 2.** Regression lines of the number of correct choices made by personality type and treatment in the 2 choice discrimination task. Figure 2A shows regression lines for control fish and 2B shows regression lines for treatment fish. The bold group is in red and the shy group is in blue. Shaded regions indicate a 95% confidence interval.
